# Multi-Cellular Human Liver Organoids Enable Complete Maturation of Induced Pluripotent Hepatocyte-like Cells Through Purely Endogenous Signals

**DOI:** 10.1101/2025.10.09.681454

**Authors:** Neeti N. Gandhi, Padmavathy Rajagopalan

## Abstract

Induced pluripotent stem cells (iPSCs) require further maturation before they can substitute primary human cells. Induced pluripotent stem cell hepatocyte-like cells (iHLCs) exhibit significantly lower hepatic functions than primary human hepatocytes (PHHs). Maturation of iHLCs has relied unsuccessfully on administering chemical cocktails in widely differing temporal patterns that are 10^6^-fold higher in concentration than *in vivo*. Hence, there is no reproducible approach for iHLC maturation. We report the assembly of a multicellular 3D human liver organoid that recapitulates the *in vivo* hepatic microenvironment. Intra-and intercellular signaling between human hepatic cells and iHLCs results in their maturation. Within seven-days, iHLCs in organoids expressed markers of hepatocyte maturation that were statistically similar to PHHs including alphafetoprotein (AFP), hepatic nuclear factor (HNF)-4𝛼, and albumin. Ki67^+^ iHLCs decreased by 2-fold from Days 1 to 14. Expression of two cytochrome P450 (CYP) enzymes, CYP3A4 and CYP2E1, in iHLCs were statistically similar to PHHs by Days 7 and 14, respectively. Biotransformation of acetaminophen and ethanol were statistically similar to PHHs by Day 14. On Day 1, the concentration of endogenously secreted prostaglandin E2 (PGE2) was identical to values reported in adult humans. Over the 14-day culture, the concentrations of endogenously secreted hepatocyte growth factor (HGF) and Oncostatin M (OSM) increased until they were 26-36% of *in vivo* values. The organoids are secreting critically important maturation molecules that are similar to levels reported in healthy humans. These trends demonstrate how closely the multi-cellular organoids are emulating i*n vivo*-like behavior while undergoing further maturation.

## 1. Introduction

Primary human cells are critical for biomedical translational science however, they are very difficult to obtain. The tissue engineering community needs a sustainable source of primary human cells which are usually obtained from biopsies or cadavers while exhibiting donor-to-donor variations (1). Induced pluripotent stem cell (iPSC)-derived hepatocyte like cells (iHLCs) are considered a potential replacement for primary human hepatocytes (PHHs) since they can be obtained non-invasively (2, 3). Although iHLCs have been used in disease modeling, drug development, and regenerative medicine (4), several drawbacks limit their potential as a replacement to PHHs (2, 3, 5).

First, many protocols that have been used to differentiate and mature iHLCs (6–16) rely on generating artificial conditions that deviate significantly from the *in vivo* hepatic environment. A common approach is the addition of chemical cocktails, including different growth factors and their agonists or chemical molecules (12, 13, 15, 16). In the body, growth factors are present in the pg-ng/mL range (17). However, many maturation protocols administer chemicals at concentrations up to 10^6^-fold higher (12, 13, 15, 16), which could potentially oversaturate receptors and alter signaling pathways and gene expression. Second, variations among publications in the temporal administration of chemical cocktails have led to a lack of a systematic, repeatable approach to mature and differentiate iHLCs *in vitro* (12–16). Other protocols include culturing the cells on polymeric scaffolds, co-culturing with hepatic and non-hepatic cells, 2D and spheroid cultures, and genetic reprogramming with the addition of various transcription factors (6–10). Third, despite the different methods developed to mature them, iHLCs continue to express fetal markers such as alphafetoprotein (AFP) and cytochrome P450 (CYP) 3A7 and even exhibit unstable phenotypes (6–16).

During liver organogenesis, intercellular signaling from neighboring endothelial, developing stellate, and hematopoietic cells can regulate and induce multiple pathways for hepatocyte progenitor cells to undergo differentiation and maturation (18, 19). Motivated by this observation, we developed a reproducible methodology that used purely endogenous signals to promote the maturation of IHLCs. Our goal was to remove the limitations of existing protocols, especially to avoid overwhelming the cells with chemicals to stimulate developmental and functional pathways. We designed a novel 3D multicellular organoid that closely mimicked the tissue microenvironment in the liver. These organoids contained iHLCs, LSECs and KCs arranged in a stratified architecture of the liver sinusoid found *in vivo*. Our goal was to demonstrate that endogenously secreted molecules drive iHLCs to maturation via signaling between the three hepatic cells (**Figure 1A**).

**Figure 1.**
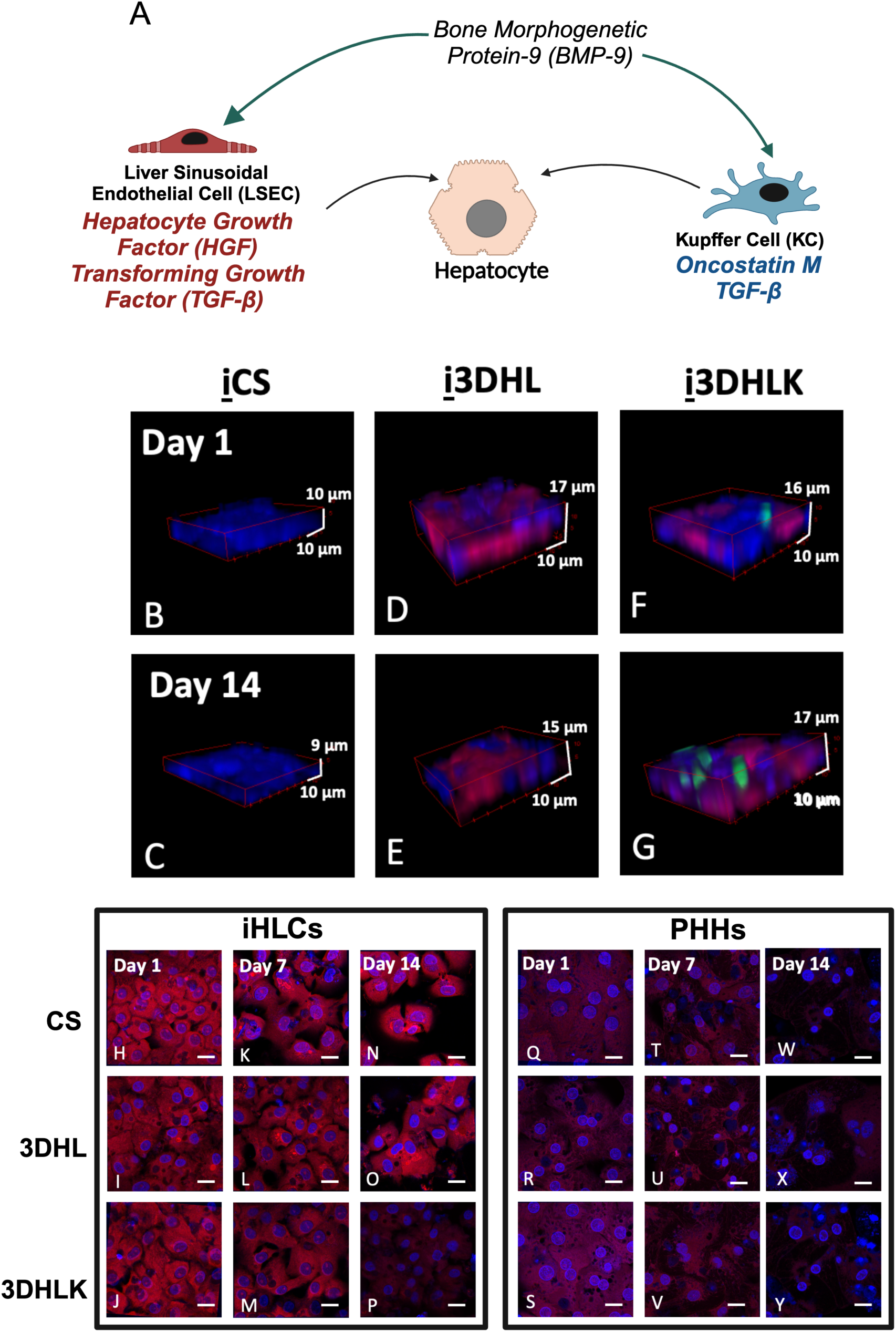
(**A**) Molecules secreted by LSECs and KCs that can signal maturation of hepatocytes during liver embryogenesis and development. 3D visual representations of **i**CS (**B**, **C**), **i**3DHL (**D**, **E**), and **i**3DHLK (**F**, **G**) cultures/ organoids on Days 1 and 14. iHLCs are immunostained for albumin (blue), LSECs for CD32b (red), and KCs for CD163 (green). Immunostaining for AFP (red) and nuclei (blue) for iHLC (**H-P**) and PHH (**Q-Y**) cultures on Days 1, 7 and 14. Scale bars = 20 𝜇m. n = 3 biological replicates. Image analysis conducted on ≥12 images with ≥ 550 cells images for each culture.

We provide a comprehensive comparison between iHLC and PHH markers to demonstrate that iHLCs in multi-cellular organoids are virtually indistinguishable from PHHs. To investigate how endogenous intercellular signaling may drive iHLC maturation, we measured the secretion of critical drivers of maturation: hepatocyte growth factor (HGF), prostaglandin E2 (PGE2), and Oncostatin M (OSM) in the organoids. We show that the secretion of PGE2 may be the key molecule to drive maturation. Since the continued expression of PGE2 may be harmful to liver cells, we also sought to elucidate how the temporal concentrations of this prostaglandin can be modulated by liver-specific proteins.

## 2. Results

When referring to the multicellular models, the notation will indicate the following. Hepatocytes, are denoted by ‘**H**’, with‘**<u>i</u>**<u>’</u> indicating iHLCs () and‘**p**’ meaning PHHs. LSECs and KCs are referred to as ‘**L**’ or ‘**K**’, respectively. Organoids are denoted by **3D**. Organoid models with iHLCs, LSECs, and KCs are referred to as **i**3DHLK, while **i**3DHL indicates organoids with iHLCs and LSECs, The counterparts that contain PHHs instead of iHLCS are referred to by **<u>p</u>**3DHLK and **<u>p</u>**3DHL. Collagen sandwich (CS) models with iHLCs or PHHs only will be referred to as **i**CS or **<u>p</u>**CS, respectively.

### 2.1 Multicellular iHLC Organoids Maintain 3D Structure

iHLC organoids were immunostained with cell-specific markers (**Supplementary**, **Figure S1**, **Table S1**). iHLCs were identified by albumin (blue), LSECs by CD32b (red), and KCs by CD163 (green) expression (**Figures 1B-G**). Throughout the 14-day culture period, the expression of albumin (iHLCs), CD32b (LSECs) and CD163 (KCs) was observed indicating that all cells maintained their phenotypic markers (**Movies S1-6)**.

Although, the heights of organoids remained constant for all iHLC models over the 14 days, there were differences between the heights of mono-and multi-cellular cultures. (**Table S2**). By Day 14, the height of **i**3DHL and **i**3DHLK organoids was up to approximately 1.6-1.8-fold higher compared to **i**CS cultures due to the inclusion of LSECs and KCs.

### 2.2 Fluorescence Intensity of AFP Decreases in Maturing iHLCs

In differentiation and maturation protocols, the expression of AFP, a fetal marker, is used to determine if stem cells have committed to a hepatic lineage (14). On Day 1, differences in AFP fluorescence intensities were statistically insignificant (*p* > 0.05) across all iHLC models (**Supplementary, Figures 1H-J**). By Days 7 and 14, the fluorescence intensity for AFP in **i**3DHLK organoids decreased (*p* < 0.05) by approximately 1.6-fold compared to **i**CS and **i**3DHL cultures (**Figures 1K-P**). There was no significant difference in AFP expression between PHH cultures at any time point (**Figures 1Q-Y**). While AFP fluorescence intensity in PHH cultures was 2.2-fold lower (*p* < 0.05) on Day 1 compared to iHLC organoids (**Figures 1H-J, Q-S**) by Days 7 and 14, there was no difference (*p* > 0.05) between **i**3DHLK and **<u>p</u>**3DHLK organoids (**Figures 1M, P, V, Y, Figures S2A-C**). This trend indicates that iHLCs, specifically in the presence of LSECs and KCs, are beginning to adopt a mature phenotype.

### 2.3 Decrease in HNF4***𝜶***^+^ Correlates to an Increase in Albumin^+^ iHLCs

HNF4α, a marker of immature hepatocytes, plays a role in regulating liver development and maturation (20). Although HNF4α is present in adult hepatocytes to maintain liver function, its expression is lower compared to hepatoblasts (21). On Day 1, approximately 50% of cells in all iHLC cultures were HNF4α-positive (HNF4α^+^) (**Supplementary, Figures 2A-C)**. By Day 7, the percent of HNF4α^+^ cells in **i**3DHLK organoids was lower (30.0 ± 1.5%, *p* < 0.05) than **i**CS cultures (45.3 ± 3.5%) (**Figures 2D-F**). By Day 14, both **i**3DHL (30.0 ± 4.5%) and **i**3DHLK (35.9 ± 1.0%) organoids exhibited a significant (*p* < 0.05) decrease in the percentage of HNF4α^+^ cells compared to **i**CS cultures (53.7 ± 3.5%) (**Figures 2G-I**). In comparison, approximately 20% of PHHs were HNF4α^+^ in all cultures over the 14-day culture (**Figures 2J-R**). By Day 7, the percentage of HNF4α^+^ cells was similar (*p* > 0.05) between **i**3DHLK and **<u>p</u>**3DHLK organoids (**Figures 2F, O**).

**Figure 2.**
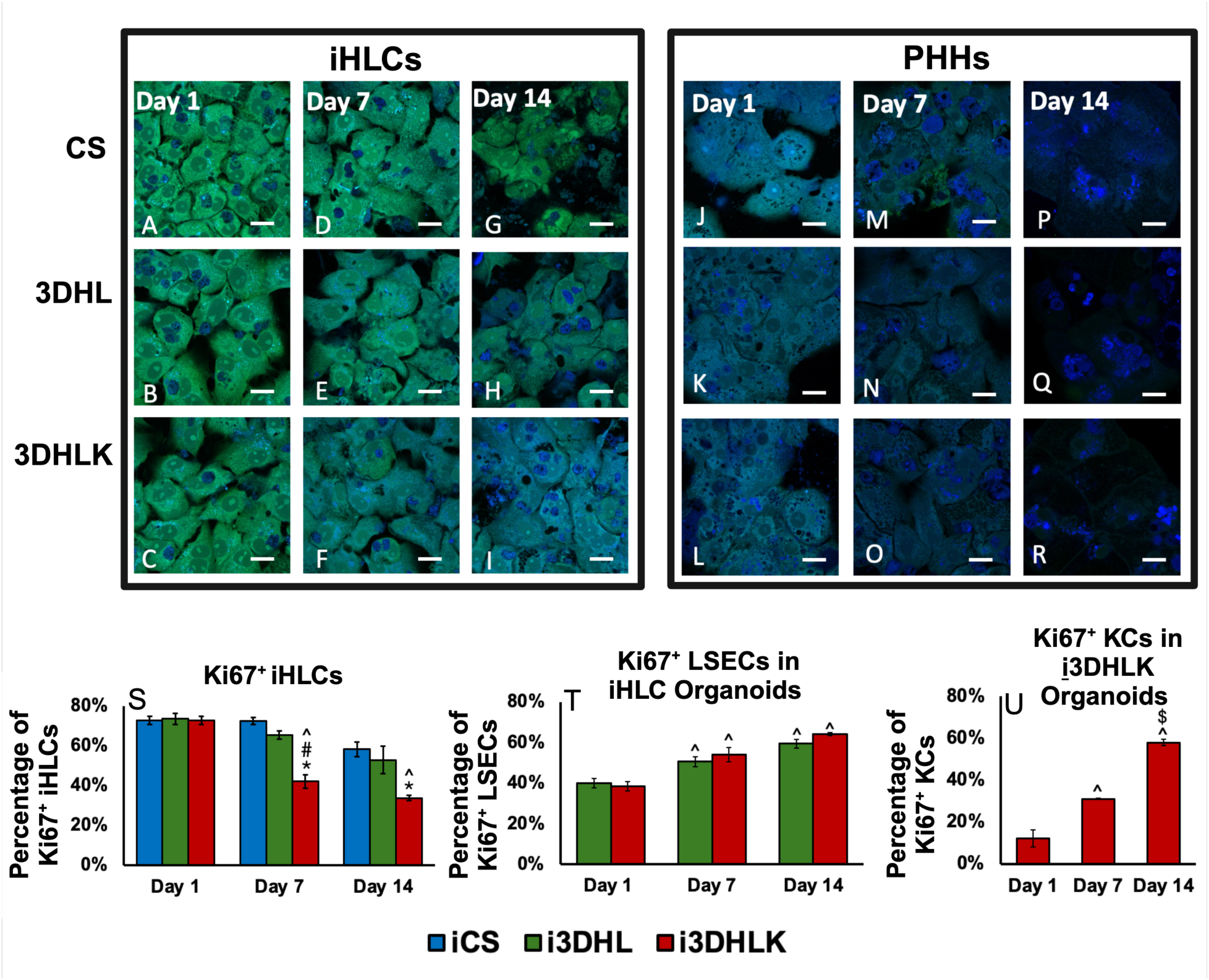
Immunostaining for HNF4𝛼 (green) and albumin (blue) in iHLC (**A-I**) and PHH (**J-R**) cultures on Days 1, 7, and 14. Scale bars = 20 𝜇m. n = 3 biological replicates. Image analysis conducted on ≥12 images with ≥ 550 cells images for each culture. Proliferating cells in iHLC organoids from Days 1 through 14. (**S**) Percentage of Ki67^+^ iHLCs on Days 1 – 14. (**T**) Percentage of Ki67^+^ LSECs in iHLC organoids on Days 1 – 14. (**U**) Percentage of Ki67^+^ KCs in iHLC organoids on Days 1 - 14. n = 3 biological replicates. Image analysis conducted on ≥12 images for each culture with ≥ 250 hepatocytes, ≥ 70 LSECs, and ≥ 100 KCs. \**p* < 0.05 indicates statistical significance compared to **i**CS culture. ^#^*p* < 0.05 indicates statistical significance compared to **i**3DHL model. ^*p* < 0.05 indicates statistical significance compared to Day 1. ^$^*p* < 0.05 indicates statistical significance compared to Day 7.

In contrast, it took up to Day 14 for the percentage of HNF4α^+^ cells in **i**3DHL and **i**3DHLK organoids to be equivalent to PHH models (*p* > 0.05) (**Figures 2H, I, Q, R, Figure S3A-C**).

Albumin secreted by adult hepatocytes (2) is used as an indicator of maturation (6, 14). We first present results on albumin-positive cells (albumin^+^) which are further validated by secretion (**Section 2.6**). On Day 1, approximately 50% of iHLCs are albumin^+^ in all cultures (**Supplementary**, **Figures 2A-C**). On Day 7, 70.0 ± 1.5% of iHLCs in **i**3DHLK organoids were albumin^+^(*p* < 0.05) compared to **i**CS cultures (54.7 ± 3.5%, **Figures 2D-F**). By Day 14, the percentage of albumin^+^ iHLCs in **i**3DHL (63.7 ± 4.5%) and **i**3DHLK (64.1 ± 1.0%) organoids was higher (*p* < 0.05) than **i**CS cultures (46.3 ± 3.5%, **Figures 2G-I**). In PHH cultures, there were no significant (*p* > 0.05) differences in albumin^+^ cells between any timepoints (**Figures 2J-R**). Interestingly, by Day 7, the percentage of albumin^+^ cells were similar (*p* > 0.05) between **i**3DHLK and **<u>p</u>**3DHLK organoids (**Figures 2F, O**). It took 14 days for the percentages of albumin^+^ iHLCs in **i**3DHL and **i**3DHLK organoids to be equivalent to PHH models (*p* > 0.05) (**Figures 2H, I, Q, R, Figure S3D-F**). These trends correlate with the decrease in HNF4𝛼 expression, suggesting that an increase in mature hepatic characteristics is accompanied by a decrease in fetal markers (20).

### 2.4 Decreases in Ki67^+^ iHLCs through Day 14 in i3DHLK organoids

Mature hepatocytes usually exist in a non-proliferative state (1). In contrast, iHLCs are known to multiply (2). To identify if there were temporal shifts, the organoids were immunostained with Ki67, a marker that indicates that a cell may be entering a proliferative state (22) (**Supplementary, Figure 2S-U, Figure S4)**.

On Day 1, approximately 70% of iHLCs were Ki67^+^ in all cultures (**Figures 2S, Figures S4A-C**). However, by Day 14, the percentage of Ki67^+^ iHLCs in **i**3DHLK organoids (33.7 ± 1.3%) had decreased by approximately 2-fold compared to Day 1 (**Figures S4G-I**). In PHH cultures, the percentage of Ki67^+^ hepatocytes remained approximately 10% (**Figures S4J-R**, **Figure S4AK**). The significant decrease in Ki67^+^ iHLCs in **i**3DHLK organoids by Day 14 demonstrates that cells are progressing towards a non-proliferative state, a characteristic of adult hepatocytes.

Hepatic non-parenchymal cells exhibited differing proliferation trends. Ki67^+^ LSECs increased approximately 10% (*p* < 0.05) at each time point in iHLC organoids (**Figures 2T, Figures S4S-X)**, Since LSECs proliferate primarily during liver regeneration and development, their proliferation was only detected in iHLC organoids (**Figure S4AL)**. KCs exhibited greater changes in proliferation in iHLC organoids. Ki67^+^ KCs in **i**3DHLK organoids increased (*p* < 0.05) approximately 4.7-fold from Day 1 (12.2 ± 4.1%) through Day 14 (57.9 ± 1.5%) (**Figures 2U, Figure S4AE-AG**) During this same period, the percentage of Ki67^+^ KCs in PHH organoids decreased (*p* < 0.05) (**Figures S4AH-AJ, Figure S4AM**). KC proliferation can occur due to the secretion of HGF, a potent mitogen (23). In iHLC organoids, HGF increased significantly from Day 1 onwards (**Section 2.6**).

In order to understand if the potential shift to a proliferative state by KCs could result in an inflammatory microenvironment, we measured the endogenous levels of TGF-β1 (**Supplementary**), since this cytokine can cause inflammation (24). TGF-β1 secretion decreased by approximately 3.3-fold in **i**3DHLK organoids from Days 1 to 14 (**Table S3**), suggesting that while KCs may be undergoing a shift towards a proliferative state, the organoids did not exhibit inflammation. TGF-β1 secretion did not change in **<u>p</u>**3DHLK organoids from Days 1 to 14, which corresponds to the virtually constant percentages of Ki67^+^ cells in PHH cultures.

### 2.5 CYP2E1 and CYP3A4 Fluorescence Intensity and Biotransformation Increase in Maturing iHLCs

PHHs are used to test CYP-mediated biotransformation of drugs, toxicants, and xenobiotics (25). iHLCs have been reported to exhibit up to 40,000-fold lower expression of CYP genes than PHHs making it difficult to substitute these cells (2, 5). We measured the fluorescence intensity and biotransformation of CYP2E1 and CYP3A4 (**Supplementary**) (26).

#### CYP2E1 Fluorescence Intensity

The fluorescence intensity of CYP2E1 was statistically insignificant (*p* > 0.05) between iHLC cultures on Days 1 and 7 (**Supplementary, Figures 3A-F**). However, by Day 14, fluorescence intensity in both **i**3DHL and **i**3DHLK organoids was higher (approximately 2-fold, *p* < 0.05) compared to **i**CS cultures (**Figures 3G-I).** There were no significant (*p* < 0.05) differences in CYP2E1 fluorescence intensity in PHH cultures on any day (**Figures 3J-R**). Compared to iHLC cultures, the fluorescence intensity of CYP2E1 was up to 1.7-fold higher (*p* < 0.05) in PHH cultures on Days 1 and 7 (**Figures 3A-F, J-O**). However, by Day 14, CYP2E1 fluorescence intensity in **i**3DHL and **i**3DHLK organoids was similar (*p* > 0.05) to **<u>p</u>**3DHL and **<u>p</u>**3DHLK organoids (**Figures 3H, I, Q, R, Figures S2D-F**).

**Figure 3.**
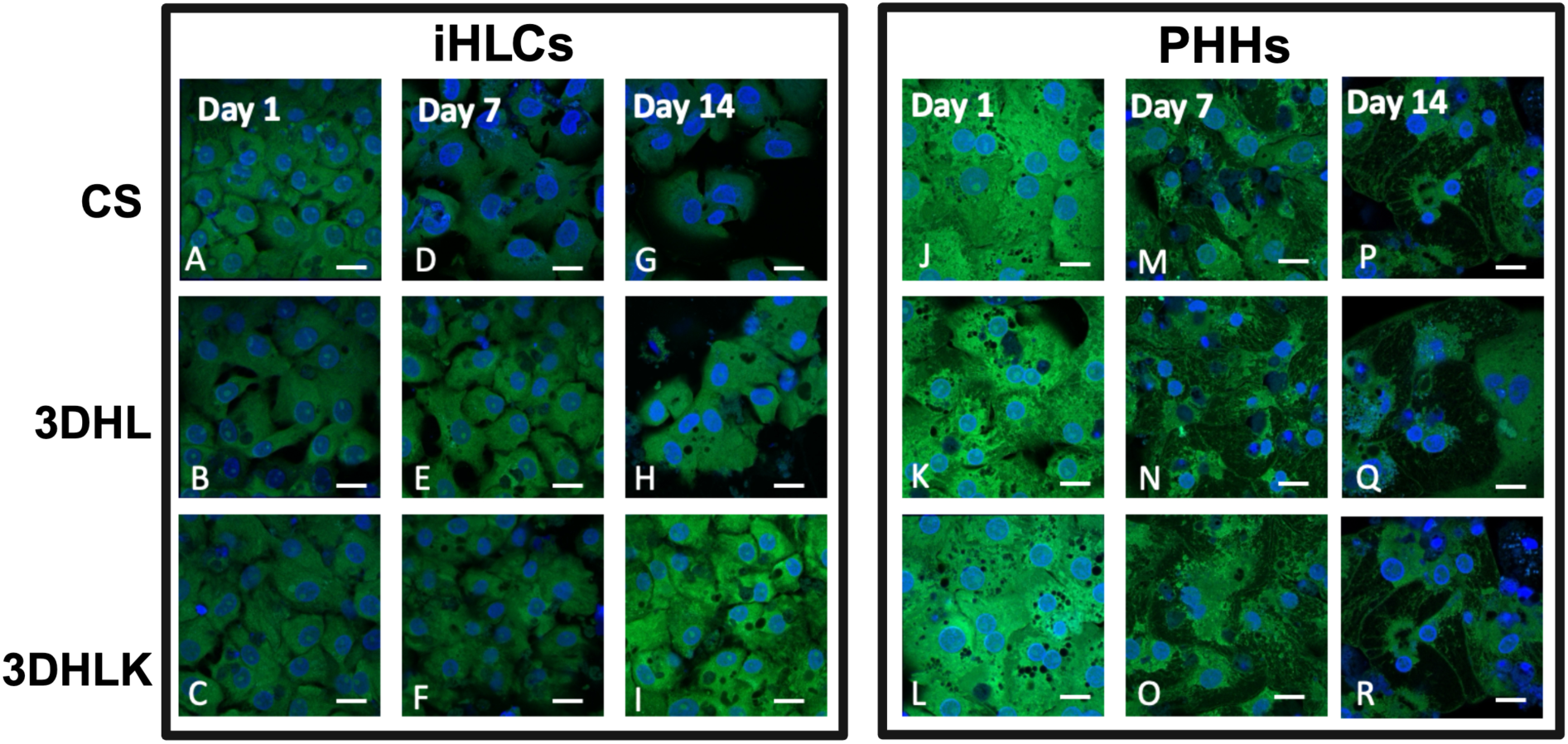
Immunostaining for CYP2E1 (green) and nuclei (blue) for iHLC (**A-I**) and PHH (**J-R**) cultures on Days 1, 7 and 14. Scale bars = 20 𝜇m. n = 3 biological replicates. Image analysis conducted on ≥12 images with ≥ 550 cells images for each culture.

#### Effects of CYP2E1 Biotransformation

APAP is the leading cause of acute liver failure in the U.S (27). EtOH can result in cirrhosis, fatty liver, and cancer (28). (**Supplementary**). When APAP (2.5 mM, LC_50_) was administered on Day 1, iHLC cultures exhibited no response to the toxicant (**Table S4**). APAP administration on Day 7 resulted in higher (*p* < 0.05) apoptosis in **i**3DHLK organoids compared to **i**CS or **i**3DHL cultures (**Table 1**). The percentage of apoptotic cells in **i**3DHLK organoids did not change (*p* > 0.05) when APAP was administered on Day 14 (**Table 1**).

**Table 1.** Percentages of apoptotic, necrotic, and total cell death measured 24 h after 2.5 mM APAP administration. n ≥ 3 biological replicates.

| Model | Administration on Day 7 |  |  | Administration on Day 14 |  |  |
| --- | --- | --- | --- | --- | --- | --- |
|  | Apoptosis (%) | Necrosis (%) | Cell Death (%) | Apoptosis (%) | Necrosis (%) | Cell Death (%) |
| iCS | $2.2 \pm 1.2$ | $6.2 \pm 3.4$ | $8.3 \pm 2.9$ | $3.9 \pm 1.7$ | $4.5 \pm 0.9$ | $8.4 \pm 2.5$ |
| i3DHL | $1.7 \pm 0.9$ | $5.0 \pm 3.2$ | $6.8 \pm 3.8$ | $2.1 \pm 3.1$ | $2.9 \pm 3.1$ | $5.0 \pm 2.7$ |
| i3DHLK | $19.6 \pm 1.6$ | $7.9 \pm 2.4$ | $27.4 \pm 0.8$ | $16.5 \pm 2.8$ | $14.1 \pm 2.1$ | $30.6 \pm 2.6$ |
| pCS | $12.6 \pm 2.2$ | $4.5 \pm 1.1$ | $17.2 \pm 3.3$ | $16.0 \pm 0.7$ | $20.6 \pm 4.2$ | $36.6 \pm 3.9$ |
| p3DHL | $6.2 \pm 1.0$ | $6.2 \pm 3.4$ | $12.4 \pm 3.7$ | $18.4 \pm 3.4$ | $22.4 \pm 6.6$ | $40.8 \pm 6.4$ |
| p3DHLK | $16.3 \pm 2.9$ | $27.3 \pm 2.8$ | $43.6 \pm 2.7$ | $18.1 \pm 1.1$ | $16.7 \pm 2.0$ | $34.8 \pm 0.8$ |

The percent of apoptotic cells was approximately 16% in both **i**3DHLK and **<u>p</u>**3DHLK organoids when APAP was added on Day 7 (**Table 1**). However, PHH models exhibited approximately 3.5-fold higher (*p* < 0.05) necrosis (26) (**Table 1**). Due to higher necrotic death in the PHH cultures, overall cell death was higher when compared to iHLC organoids.

When EtOH (160 mM, LC_50_) was administered on Day 1, iHLC organoids exhibited no response to this toxicant (**Table S6**). By Day 7, apoptosis was higher (*p* < 0.05) in the **i**3DHLK organoids compared to **i**3DHL or **i**CS cultures while necrosis was similar (*p* > 0.05) in all models (**Table 2**).

**Table 2.** Percentages of apoptotic, necrotic, and total cell death measured 24 h after 160 mM EtOH administration. n ≥ 3 biological replicates.

| Model | Administration on Day 7 |  |  | Administration on Day 14 |  |  |
| --- | --- | --- | --- | --- | --- | --- |
|  | Apoptosis (%) | Necrosis (%) | Cell Death (%) | Apoptosis (%) | Necrosis (%) | Cell Death (%) |
| iCS | $1.8 \pm 1.4$ | $3.6 \pm 1.4$ | $5.3 \pm 2.2$ | $12.3 \pm 0.4$ | $4.2 \pm 1.1$ | $16.4 \pm 0.8$ |
| i3DHL | $3.8 \pm 1.0$ | $3.5 \pm 1.1$ | $7.3 \pm 2.0$ | $3.6 \pm 2.0$ | $6.5 \pm 0.2$ | $10.1 \pm 2.0$ |
| i3DHLK | $20.6 \pm 1.9$ | $9.8 \pm 2.9$ | $30.3 \pm 1.9$ | $24.7 \pm 1.12$ | $5.5 \pm 1.6$ | $30.2 \pm 2.1$ |
| pCS | $12.7 \pm 1.3$ | $6.6 \pm 1.2$ | $19.3 \pm 1.5$ | $22.1 \pm 3.5$ | $25.7 \pm 4.2$ | $47.9 \pm 6.8$ |
| p3DHL | $5.3 \pm 1.1$ | $6.0 \pm 3.4$ | $11.3 \pm 4.4$ | $21.3 \pm 7.0$ | $24.3 \pm 7.1$ | $45.6 \pm 13.1$ |
| p3DHLK | $19.7 \pm 1.0$ | $29.0 \pm 3.2$ | $48.6 \pm 4.2$ | $21.6 \pm 7.3$ | $36.0 \pm 2.7$ | $57.5 \pm 7.9$ |

When EtOH was added either on Day 7 or 14, both **i**3DHLK and **<u>p</u>**3DHLK organoids exhibited similar (*p* > 0.05) percentages of apoptotic cells (**Table 2**). However, the percentage of necrotic cells in **<u>p</u>**3DHLK organoids was consistently higher (*p* < 0.05) than **i**3DHLK models. iHLC and PHH organoids exhibited similar levels of apoptosis but differed in necrotic cell death due to EtOH administration.

#### CYP3A4 Fluorescence Intensity

CYP3A4 is an enzyme that metabolizes approximately 50% of drugs and pharmaceuticals (26). The fluorescence intensity of CYP3A4 was identical in all iHLC models on Day 1 (**Supplementary**, **Figures 4A-C**). The fluorescence intensity of CYP3A4 was 2.1-fold and 2.8-fold higher (*p* < 0.05) in **i**3DHLK cultures on Days 7 and 14, respectively compared to **i**CS cultures (**Figures 4D, F, G, I**). The fluorescence intensity did not vary significantly in any PHH culture at any timepoint **(Figures 4J-R**). On Days 7 and 14, CYP3A4 fluorescence intensity in **i**3DHLK and **<u>p</u>**3DHLK organoids was similar (*p* > 0.05) (**Figures S2G-I**), indicating that iHLCs were now expressing another marker of mature hepatocytes.

**Figure 4.**
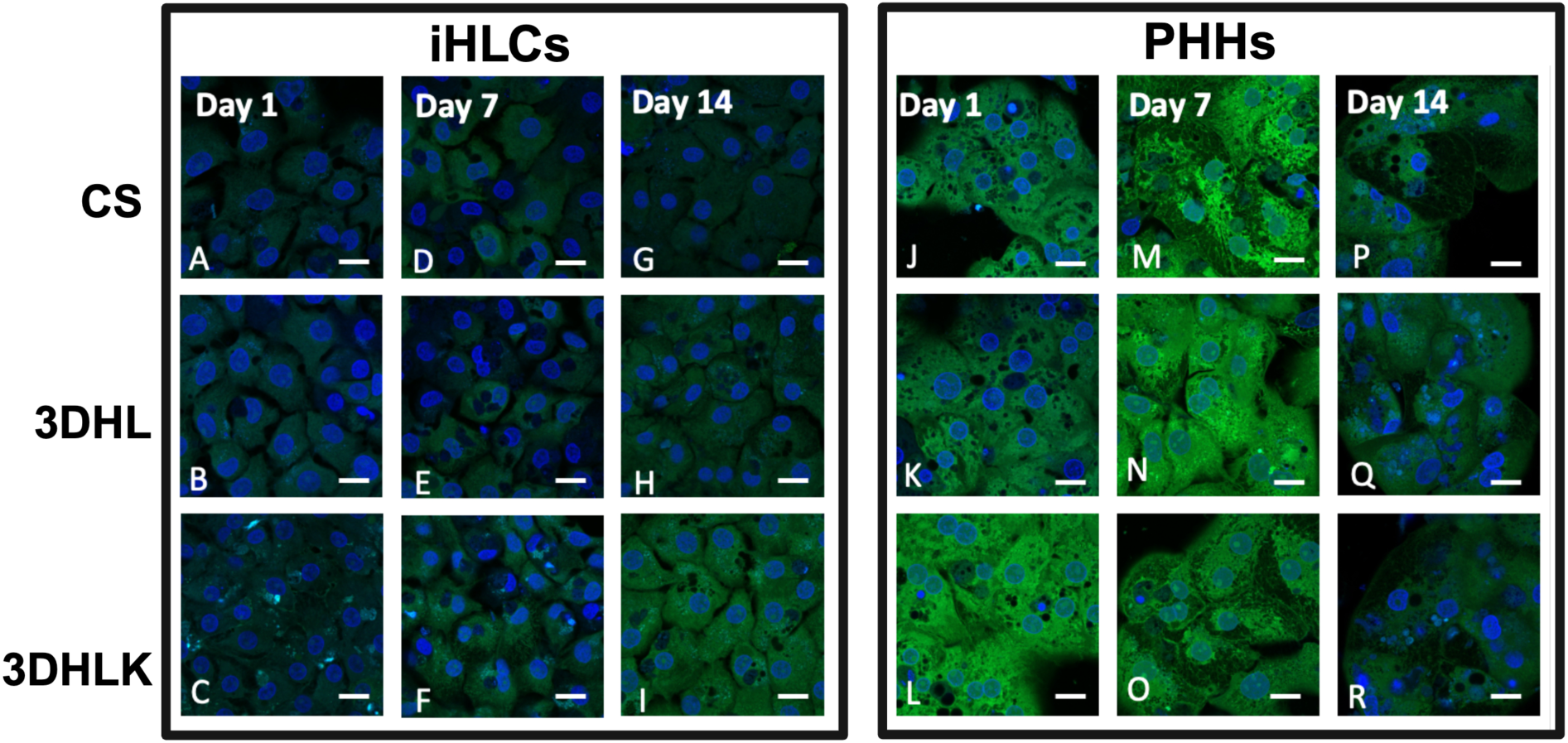
Immunostaining for CYP3A4 (green) and nuclei (blue) for iHLC (**A-I**) and PHH (**J-R**) cultures on Days 1, 7 and 14. Scale bars = 20 𝜇m. n = 3 biological replicates. Image analysis conducted on ≥12 images with ≥ 550 cells images for each culture.

#### Effects of CYP3A4 Biotransformation

When RIF, an anti-tuberculosis antibiotic (29) (0.36 𝜇M, EC_50_) (30) was administered on Day 1, iHLC organoids did not exhibit any response (*p* > 0.05) to the toxicant (**Supplementary**, **Table S8**). After administering on either Day 7 or 14, **i**3DHLK organoids exhibited higher (*p* < 0.05) apoptosis compared to both **i**CS and **i**3DHL cultures (**Table 3**). Overall cell death was higher (*p* < 0.05) in **i**3DHLK organoids compared to **i**CS and **i**3DHL cultures on Days 7 and 14 (**Table 3**).

**Table 3.** Percentages of apoptotic, necrotic, and total cell death measured 24 h after 0.36 𝜇M RIF administration. n ≥ 3 biological replicates.

| Model | Administration on Day 7 |  |  | Administration on Day 14 |  |  |
| --- | --- | --- | --- | --- | --- | --- |
|  | Apoptosis (%) | Necrosis (%) | Cell Death (%) | Apoptosis (%) | Necrosis (%) | Cell Death (%) |
| iCS | $8.0 \pm 3.3$ | $1.8 \pm 1.4$ | $9.8 \pm 1.9$ | $3.2 \pm 0.0$ | $3.9 \pm 0.8$ | $7.1 \pm 0.8$ |
| i3DHL | $7.5 \pm 2.3$ | $1.7 \pm 1.2$ | $9.1 \pm 3.2$ | $5.6 \pm 1.5$ | $5.3 \pm 2.5$ | $10.9 \pm 3.9$ |
| i3DHLK | $20.3 \pm 2.8$ | $9.4 \pm 2.5$ | $29.6 \pm 4.9$ | $18.1 \pm 2.3$ | $14.5 \pm 2.8$ | $32.5 \pm 4.2$ |
| pCS | $9.2 \pm 2.1$ | $2.0 \pm 0.5$ | $11.2 \pm 2.5$ | $11.5 \pm 1.0$ | $2.9 \pm 0.3$ | $14.4 \pm 1.3$ |
| p3DHL | $16.8 \pm 3.0$ | $6.4 \pm 0.6$ | $23.3 \pm 2.4$ | $9.4 \pm 1.4$ | $7.6 \pm 1.1$ | $17.0 \pm 2.4$ |
| p3DHLK | $28.2 \pm 1.9$ | $15.7 \pm 2.0$ | $41.8 \pm 3.9$ | $23.5 \pm 3.1$ | $21.7 \pm 3.0$ | $45.2 \pm 1.1$ |

By Day 7, approximately 20% of cells were apoptotic in **i**3DHLK and **<u>p</u>**3DHLK (*p* > 0.05) organoids. Overall, on Day 7, cell death was higher (*p* < 0.05) in **<u>p</u>**3DHLK cultures compared to **i**3DHLK models due to higher necrosis (*p* < 0.05). However, by Day 14, there were no significant differences (*p* > 0.05) in the percentage of apoptotic or necrotic cells between **i**3DHLK and **<u>p</u>**3DHLK organoids (**Table 3**). The progressive capability in biotransformation highlights that if iHLCs are present in a microenvironment that includes LSECs and KCs, after 14 days, they can exhibit similar CYP3A4 metabolism as PHHs. We hypothesized that autocrine and paracrine signaling among the three cell types may be essential for iHLC maturation. Accordingly, we measured the concentrations of signaling molecules over the 14-day period to obtain a deeper understanding on maturation.

### 2.6 Endogenous Secretion of Maturation Molecules Promotes iHLC Maturation

We measured the endogenous secretion of HGF, OSM, and PGE2 (**Supplementary**), three molecules that are critical regulators of liver maturation (23, 31, 32), to elucidate the mechanisms that drive the decrease in fetal and increase in adult hepatocyte markers in the organoids. The secretion of each molecule was first measured in monocultures to determine which cell type was primarily responsible for its production. Endogenous secretion of proteins from NPC monocultures were only measured on Day 1 since these cells de-differentiate in 2D cultures (33).

#### LSECs Secrete HGF

In LSEC monocultures, HGF secretion was approximately 142-fold higher compared to other cell types (**Table 4**). Therefore, we identified LSECs as primarily responsible for endogenous HGF in the organoids (34). Over the 14-day culture period, HGF increased by approximately 3.1 and 2-fold in **i**3DHL and **i**3DHLK organoids, respectively (**Figure 5A, Table S10**). In **i**CS cultures, HGF remained virtually constant. By Day 14, the endogenous secretion of HGF (after subtracting culture medium contributions) reached approximately 26% (**i**3DHLK) and 36% (**i**<u>3DHL)</u> of values reported *in vivo* for adult humans (270 pg/mL) (17). In contrast, PHH organoids did not exhibit any HGF secretion (**Table 4**).

**Figure 5.**
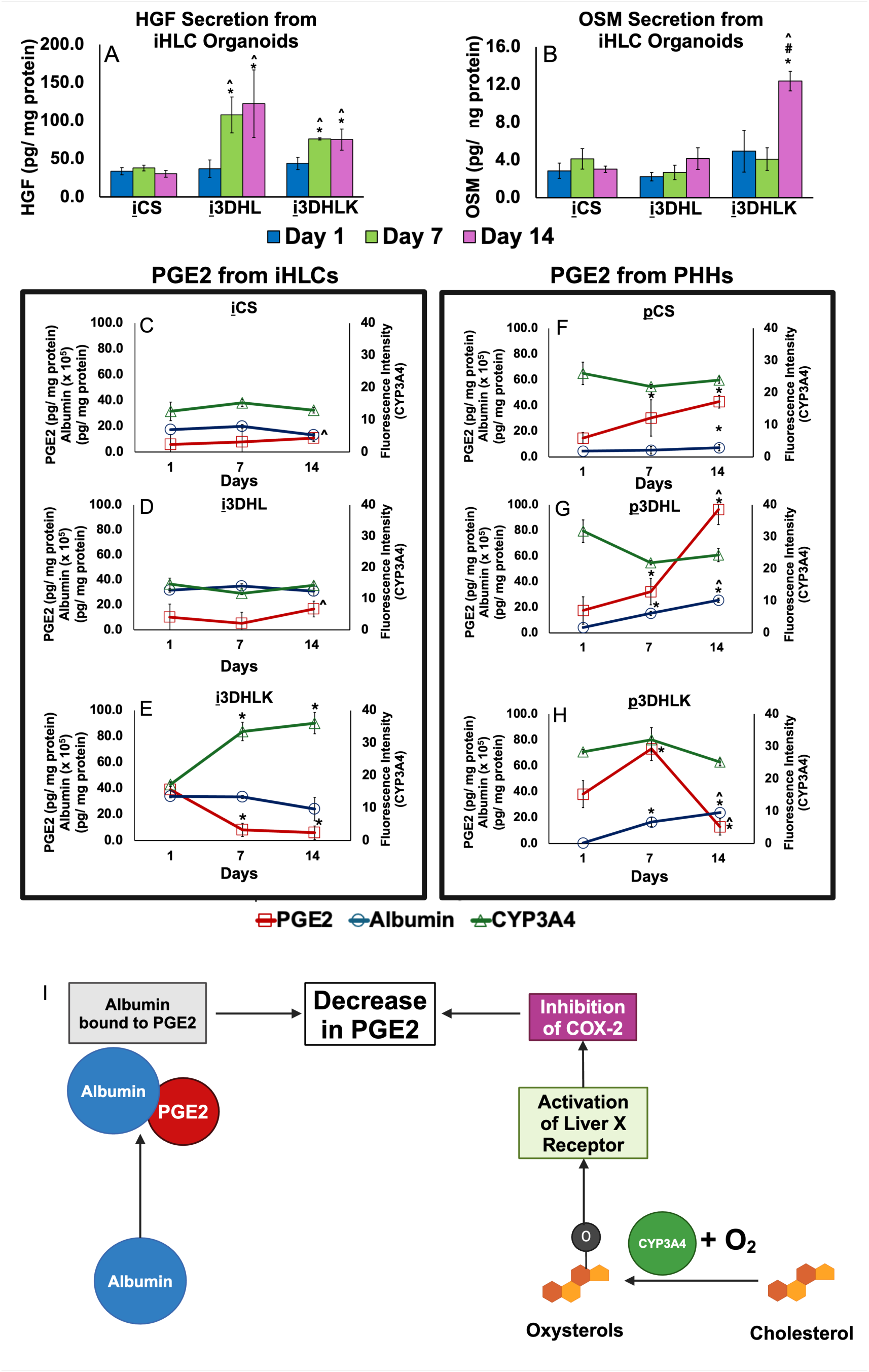
Endogenous secretion of maturation molecules in iHLC and PHH cultures. (**A**) HGF and (**B**) OSM secretion in iHLC organoids (pg/ mg protein). n = 3 biological replicates. The value obtained from culture medium was subtracted from the raw data and then normalized to protein content in the organoid at each time point. \**p* < 0.05 indicates statistical significance compared to CS model. ^#^*p* < 0.05 indicates statistical significance compared to 3DHL model. ^*p* < 0.05 indicates statistical significance compared to Day 1. PGE2 (pg/ mg protein) and albumin (x 10^5^ pg/ mg protein) secretion and fluorescence intensity (CYP3A4) from (**C-E**) iHLC and (**F-H**) PHH models. n = 3 biological replicates for each culture for all measurements. Culture medium values were subtracted from the raw data and then the concentration was normalized to protein content in the organoid at each time point. Image analysis was conducted on ≥12 images with ≥ 550 cells for each culture/organoid. \**p* < 0.05 indicates statistical significance compared to Day 1. ^^^*p* < 0.05 indicates statistical significance compared to Day 7. (**I)** On Days 7 and 14, albumin and CYP3A4 may decrease PGE2 expression. As albumin increases, it binds to and reduces the amount of available PGE2. CYP3A4 can oxidize cholesterol to form oxysterols, which activate LXR. COX-2 is inhibited when LXR is activated, leading to a decrease in PGE2.

**Table 4.** Secretion of HGF, OSM and PGE2 in LSEC, KC, iHLC, and PHH organoids/ cultures (pg/ mL). Concentrations were calculated by subtracting individual culture medium values at each time point. n = 3 biological replicates. (-) indicates that the concentration of the molecule in the culture was lower than that of the culture medium.

|  |  | HGF<br>(pg/ mL) | OSM<br>(pg/ mL) | PGE2<br>(pg/ mL) |
| --- | --- | --- | --- | --- |
| Day 1 | LSECs | 188.9 ± 14.2 | 0.2 ± 0.4 | 9.4 ± 0.5 |
|  | KCs | 1.3 ± 1.2 | 7.8 ± 2.8 | 4.2 ± 3.7 |
|  | iCS | 29.1 ± 4.0 | 3.0 ± 1.3 | 5.0 ± 6.7 |
|  | i3DHL | 31.6 ± 9.7 | 2.7 ± 1.2 | 8.7 ± 8.8 |
|  | i3DHLK | 35.0 ± 6.3 | 3.9 ± 1.8 | 31.0 ± 4.9 |
|  | pCS | – | – | 7.5 ± 2.2 |
|  | p3DHL | – | – | 10.1 ± 6.2 |
|  | p3DHLK | – | – | 20.4 ± 11.0 |
| Day 7 | iCS | 32.2 ± 5.3 | 3.3 ± 0.9 | 6.3 ± 11.0 |
|  | i3DHL | 85.7 ± 26.1 | 2.0 ± 0.6 | 3.9 ± 6.7 |
|  | i3DHLK | 65.2 ± 1.3 | 3.5 ± 1.0 | 6.9 ± 4.54 |
|  | pCS | – | – | 16.7 ± 7.8 |
|  | p3DHL | – | – | 18.4 ± 5.9 |
|  | p3DHLK | – | – | 43.1 ± 1.3 |
| Day 14 | iCS | 33.4 ± 6.0 | 3.4 ± 0.2 | 10.9 ± 2.7 |
|  | i3DHL | 99.2 ± 37.7 | 3.4 ± 0.9 | 13.7 ± 5.1 |
|  | i3DHLK | 69.7 ± 12.7 | 11.4 ± 1.0 | 5.6 ± 4.9 |
|  | pCS | – | – | 25.3 ± 10.9 |
|  | p3DHL | – | – | 38.3 ± 5.4 |
|  | p3DHLK | – | – | 5.4 ± 4.7 |

#### KCs Secrete OSM

In monocultures, OSM secretion by KCs was up to 35.4-fold and 2.6-fold higher compared to LSECs and iHLCs on Day 1, respectively **(Table 4**) (19). OSM secretion exhibited a temporal increase of 2.9-fold over 14 days only in **i**3DHLK organoids (**Figure 5B, Table S11**). Once again, by Day 14, the concentration of OSM in **i**3DHLK organoids was approximately 32% of values observed i*n vivo* (35.8 pg/mL) in adult humans (**Table 4**). The concentrations of HGF and OSM secreted by iHLC organoids approached values observed in humans suggesting that the cultures are able to drive maturation even at these low levels.

On Day 14, OSM secretion was higher (*p* < 0.05) than Days 1 or 7 only in **i**3DHLK organoids (**Figure 5B)**. The concentration of OSM at earlier time-points is only about 10% of the value reported i*n vivo*. We hypothesize that the presence of Ki67^+^ KCs are contributing to the increase in OSM in order to promote maturation mechanisms. In PHH organoids, endogenous OSM was not detected throughout the culture period (**Table 4**, **Table S11**).

#### iHLC and PHH Organoids Exhibit Different Temporal Patterns for PGE2 Secretion

Although all cells secreted PGE2, LSEC and PHH monocultures exhibited approximately 2-fold higher secretion compared to KCs and iHLCs (**Table 4**). In iHLC organoids, we focus upon the tri-cellular systems since they exhibit the greatest maturation and most significant temporal variations in PGE2 levels. On Day 1, the PGE2 concentration in **i**3DHLK organoids was 31 ± 4.9 pg/mL, which is very close to the *in vivo* value of 24.2 pg/mL (**Table 4**, **Figure 5E**). By Days 7 and 14, PGE2 levels went down by 4.9-fold and 6.5-fold, respectively (**Discussion**, **Table S12**). During the culture period, Ki67^+^ iHLCs decreased by approximately 2-fold, which correlated with the decrease in PGE2. Since PGE2 is implicated in liver developmental pathways (32), its levels may be decreasing since iHLCs have begun to stop proliferating as well as exhibiting maturation markers. While **i**3DHLK organoids showed a decrease in endogenously secreted PGE2 the reverse trend was seen in i3DHL cultures suggesting that this prostaglandin may be needed in order to promote maturation due to the lack of KCs.

#### PGE2 Levels Correlate to Albumin Secretion and CYP3A4 Expression in 3DHLK Organoids

PGE2 can be modulated by the bioavailability of albumin and through the activation of the liver X receptor (LXR) (**Figure 5I**) (35, 36). CYP450 enzymes, specifically, CYP3A4, can form oxysterols that induce LXR, which in turn inhibits PGE2 (37, 38). Albumin can also modulate PGE2 bioavailability upon binding to the prostaglandin (35).

A significant increase in the fluorescence intensity of CYP3A4 was observed only in **i**3DHLK organoids on Day 7 and maintained up to Day 14 (**Section 2.5**). PGE2 secretion and CYP3A4 expression exhibit an inverse relationship in **i**3DHLK organoids throughout the culture period suggesting that CYP3A4 may be one mechanism by which PGE2 levels are modulated. However, albumin levels in iHLCs organoids did not vary significantly over time in any of the culture conditions, indicating that the bioavailability of this protein may not play a role in PGE2 regulation in **i**3DHLK organoids.

In **<u>p</u>**3DHLK organoids, albumin secretion increased approximately 4 and 5-fold on Days 7 and 14, respectively, compared to Day 1. The increase in albumin levels (**Table S13**) correlated to a 5.7-fold decrease in PGE2 by Day 14 compared to Day 7. The fluorescence intensity for CYP3A4 did not change in PHH models, suggesting that regulation of PGE2 in the presence of PHHs may be controlled by albumin bioavailability. We hypothesize that either CYP3A4 expression or albumin may contribute to the reduction in PGE2 levels but only in the presence of KCs. Taken together, we demonstrate that LSECs and KCs are primarily responsible for the secretion and regulation of molecules that drive maturation in multicellular iHLC organoids.

## 3. Discussion

The primary goal of this study was to demonstrate that the unique hepatic microenvironment established when human LSECs and KCs are in close proximity to iHLCs in an organoid can lead to complete maturation through the endogenous secretion of signaling molecules. Previous studies on iHLC maturation have reported the administration of chemical cocktails using a variety of temporal profiles where the concentrations of exogenously added reagents can be up to 6-orders of magnitude higher than present *in vivo* (12, 13, 15, 16). Non-physiological concentrations of exogenous chemicals cause alterations in cellular signaling. We demonstrate that the *in vivo*-like concentrations of endogenously secreted HGF, OSM, and PGE2 (**Table 4**) are sufficient to completely and reproducibly mature iHLCs. In the presence of both LSECs and KCs, iHLCs can mature more rapidly due to a combination of inter-and intra-cellular signaling from both hepatic cell types while the inclusion of LSECs alone requires more time for iHLC maturation.

Two critical signaling molecules that mark hepatocyte differentiation, HGF and OSM, are detected only in iHLC organoids. Even more significant was that the concentrations of each signaling molecule were similar or almost identical to those reported *in vivo* (**Table 4**) (17, 39, 40). The endogenous secretions of HGF, PGE2, and OSM signaling can initiate different pathways that either inhibit fetal or induce adult hepatic markers. The synergism between these molecules may explain maturation is more rapid in organoids when both LSECs and KCs present.

HGF, a potent mitogen, activates the c-met receptor in hepatocytes by tyrosine phosphorylation (23). LSECs may secrete this molecule to stimulate proliferation (34). HGF can also be internalized and degraded rapidly through its receptor, c-met, in hepatocytes within 30 minutes (41). This could explain why HGF concentrations are lower in organoids (**Table S10**) compared to LSEC monocultures.

PGE2 is generated through the metabolism of arachidonic acid by cyclooxygenase-2 (COX-2) (42). HGF can upregulate COX-2 expression leading to PGE2 production (43). A positive paracrine feedback mechanism occurs wherein PGE2 can induce HGF (23, 32, 43). PGE2 can also initiate the secretion of OSM from KCs (44, 45). OSM, a member of the IL-6 family, induces hepatocyte maturation during liver organogenesis (31). We hypothesize that the significant increase in OSM secretion in **i**3DHLK organoids on Day 14 is due to potential KC proliferation (**Section 2.4**).

Temporal variations in PGE2 levels were observed in PHH and iHLC organoids although their patterns differed. To understand the underlying reasons behind the fluctuation, we correlated the temporal variation of PGE2 to albumin bioavailability and CYP3A4 expression (35, 36). Albumin decreases circulating PGE2 by reducing its binding capacity to EP-receptors (35). Additionally, the decrease in PGE2 levels in **<u>p</u>**3DHLK cultures may be due to IL-4 secretion by KCs that can suppress COX-2 (46). CYP450 enzymes can form oxysterols that activate the transcriptional activity of LXR (36–38). Oxysterols have important roles in embryonic development and organ regeneration by binding to the Hedgehog signaling pathway (47), which may explain why PGE2 and CYP3A4 have opposing trends over time in **i**3DHLK organoids.

Adult hepatocytes are typically quiescent and enter the G1 phase of the cell cycle upon injury or regeneration (1). The percentage of Ki67^+^ iHLCs decreases only in **i**3DHLK organoids, indicating that the cells are transitioning from a stem-cell state to a differentiated phenotype. This trend coincides with a decrease in PGE2 levels. Continued secretion of PGE2 is detected primarily during stem cell proliferation (48). Both LSECs and KCs can proliferate during liver regeneration when mitogens such as HGF are secreted, which may explain why these NPCs only proliferate in iHLC organoids (34). Although KCs can proliferate, the decrease in TGF-β1 secretion in **i**3DHLK organoids indicates that inflammation is not occurring (24).

Cell-cell signaling has been shown to improve the functionality of iHLCs (49, 50). Takebe et. al. investigated how signaling from human umbilical vein endothelial cells and mesenchymal stem cells further differentiated iPSC-hepatic endoderm cells by generating functional and vascularized human liver buds that were transplanted in mice (49). Although the liver buds exhibited improved functionality compared to iHLCs alone, an *in vivo* environment was found to be necessary for maturation. Ouchi et. al. assembled an organoid with iHLCs, Kupffer-like cells, and hepatic stellate-like cells to recapitulate steatohepatitis (50). While the organoids were able to emulate aspects of steatohepatitis, they exhibited significant downregulation of hepatocyte genes compared to the human liver (50). Neither study reported a phenotypic or functional comparison to PHHs *in vitro* (49, 50). If PHHs are to be substituted with iHLCs *in vitro*, it is critical to elucidate how iHLCs match adult hepatocyte properties (5).

## 4. Conclusions and Future Work

The maturation of iHLCs reported herein is based on the importance of and inherent need for cell-cell communications while also providing a novel and systematic platform for liver research.

This approach relies on mimicking a hepatic microenvironment and modulated signaling instead of external additives. In the future, the inclusion of stellate cells and cholangiocytes may provide additional maturation mechanisms.

## 5. Methods

Phosphate buffered saline (PBS, 10 X), RPMI 1640, B-27, gentamicin, RIF, penicillin-streptomycin (Pen-Strep), sodium borohydride (NaBH_4_), Triton-X, bovine serum albumin (BSA), GlutaMAX, goat serum, and horse serum were obtained from Thermofisher Scientific (Waltham, MA). Glutaraldehyde, human recombinant insulin, HEPES, APA) and EtOH were obtained from Sigma-Aldrich (St. Louis, MO). Dexamethasone was obtained from MP Biomedicals (Santa Ana, CA). Fibronectin was obtained from R&D Systems (Minneapolis, MN).

### 5.1 Culturing Hepatic Cells

All cells were maintained at 37℃ in a humidified environment at 5% CO_2_. Spent culture medium was collected and stored at-80℃ until analyzed.

**<u>iHLCs</u>:** iHLCs (iCell Hepatocytes 2.0: Fujifilm (Santa Ana, CA) were seeded on collagen gels at 300,000 cells/ cm^2^ according to the manufacturer’s protocol (5) (**Supplementary**). Culture media was changed every 24 h. iHLCs were cultured for 7 days prior to organoid assembly (5).

**PHHs:** PHHs (Sekisui Xenotech, Kansas City, KS) were seeded on collagen gels at 35,000 cells/ well in a 96-well plate (26) and cultured according to the manufacturer’s protocol. Briefly, cells were thawed in OptiThaw, then seeded in OptiPlate, and maintained with media changes every 24 h thereafter with OptiCulture. Cells were cultured for 24 h prior to organoid assembly.

**<u>LSECs</u>:** LSECs (ScienCell, Carlsbad, CA) with a passage number ≤ 5 were maintained in the manufacturer’s endothelial cell medium prior to organoid assembly (26). LSEC monocultures were seeded on collagen gels on the day of organoid assembly and maintained in iHLC media.

**KCs:** KCs (Thermofisher Scientific, Waltham, MA) or Novabiosis, Durham, NC) were thawed according to the manufacturer’s protocol 24 h before organoid assembly (**Supplementary**).

### 5.2 Organoid Assembly

LSECs and KCs were mixed in a 1.1 mg/ mL collagen Type 1 solution that contained fibronectin (1% v/v). Initial seeding ratios were 5:1 hepatocytes: LSECs and 10:1 hepatocytes: KCs, which is similar to cellular ratios *in vivo* (25). To assemble the collagen sandwich (CS) models, a second layer of collagen was added on top of the iHLCs or PHHs (**Figure S1A**). All organoids were maintained for up to 14 days. Cultures were ended on Days 1, 7, or 14 post-organoid assembly (**Figure S1B**).

### 5.3 Statistical Analysis

A two-tailed Student’s t-test with 𝛼 = 0.05 was used to determine statistical significance while assuming unequal variance. The Bonferroni correction (multiple hypothesis testing) was also applied. All data are reported as the mean ± standard deviation where *n* represents the sample size.

## AUTHORS’ CONTRIBUTIONS

N.G and P.R. designed the scope of the paper. N.G. performed the experiments, and statistical analysis. N.G. and P.R. interpreted the data and wrote the manuscript.

## ACKNOWLEDGEMENTS

The authors gratefully acknowledge support for this work from the National Science Foundation (NSF 2200045, P. R.), Institute for Critical Technologies and Applied Science, Virginia Tech (P.R.) and the Computational Tissue Engineering Interdisciplinary Graduate Education Program, Virginia Tech (N. G. and P.R.). Figures 1A, 5I, and S1 were created using Biorender.com.

## Supplementary Information

### Methods

#### iHLC Culture

Plating media consisted of RPMI 1640 supplemented with B-27 (2% v/v), OSM (10 𝜇g/ mL), dexamethasone (5 mM), gentamicin (50 ng/ mL), and iCell Hepatocyte 2.0 medium supplement (2% v/v). Maintenance media consisted of RPMI 1640 supplemented with B-27 (2% v/v), dexamethasone (5 mM), and gentamicin (50 ng/ mL), and iCell Hepatocyte 2.0 medium supplement (2% v/v). The cells were initially thawed and maintained for five days in plating media and in maintenance media thereafter.

#### KC Culture

KCs from Thermofisher Scientific were maintained in Advanced DMEM supplemented with FBS (5% v/v) and a Thawing Cocktail (3.6% v/v) which comprised of Pen-Strep (10,000 U/ mL), human recombinant insulin (4 mg/ mL), GlutaMAX (200 nm), and HEPES solution (1 M) at a pH of 7.4. KCs from Novabiosis were thawed in Kupffer Plating Medium and then Kupffer Maintenance Medium from the manufacturer 24 h before organoid assembly. KC monocultures were seeded on collagen gels maintained in iHLC media.

#### Immunostaining for Phenotypic Studies

Glass-bottom well plates were activated to promote the adhesion of collagen gels using previously reported procedures (51). Cultures were fixed with a 2% (w/v) glutaraldehyde in PBS (1 X) solution followed by a 0.1% NaBH_4_ solution in PBS (1 X) and 0.1% Triton-X solution in PBS (1 X). A blocking solution of 1.5% (v/v) goat serum in 1% (w/v) BSA in PBS or 5% (v/v) horse serum in 1% (w/v) BSA in PBS (1 X) solution was added to cultures. A list of primary and secondary antibodies is presented in **Table S1**. The nuclei were stained with Hoechst 33258 (Thermofisher). Imaging was conducted on Zeiss LSM 800 and 880 confocal microscopes. Image analysis was conducted with Nikon® NIS-Elements Software. 3D renderings of the cultures/ organoids were constructed with ImageJ. Fluorescence intensity was measured for ≥ 550 cells for every culture and time point.

#### Administration of Toxicants and Measuring Mode of Cell Death

APAP (2.5 mM), EtOH (160 mM), and RIF (0.36 𝜇M) were dissolved in culture medium at either their respective LC_50_ (lethal concentration 50) or EC_50_ (half maximal effective concentration) values and administered on Days 1, 7, and 14 (26, 30, 52). Percentages of apoptotic, necrotic, and live cells were determined through a commercially available kit (Apoptotic, Necrotic, and Healthy Cell Quantification Kit, Biotium, Fremont, CA). Imaging was immediately conducted on Zeiss LSM 800 or 880 confocal microscopes. Image analysis was conducted with Nikon® NIS-Elements Software.

#### Measuring Concentrations of Signaling Molecules

The concentrations of HGF, OSM, PGE2, and albumin were measured using commercially available kits (Abcam, Cambridge, UK) as per the manufacturer’s protocol. The concentration of TGF-β1 was measured using a commercially available kit (Thermofisher). The concentrations of each protein were determined by calibrating to a standard curve at an absorbance of 450 nm.

---

**Figure S1.**
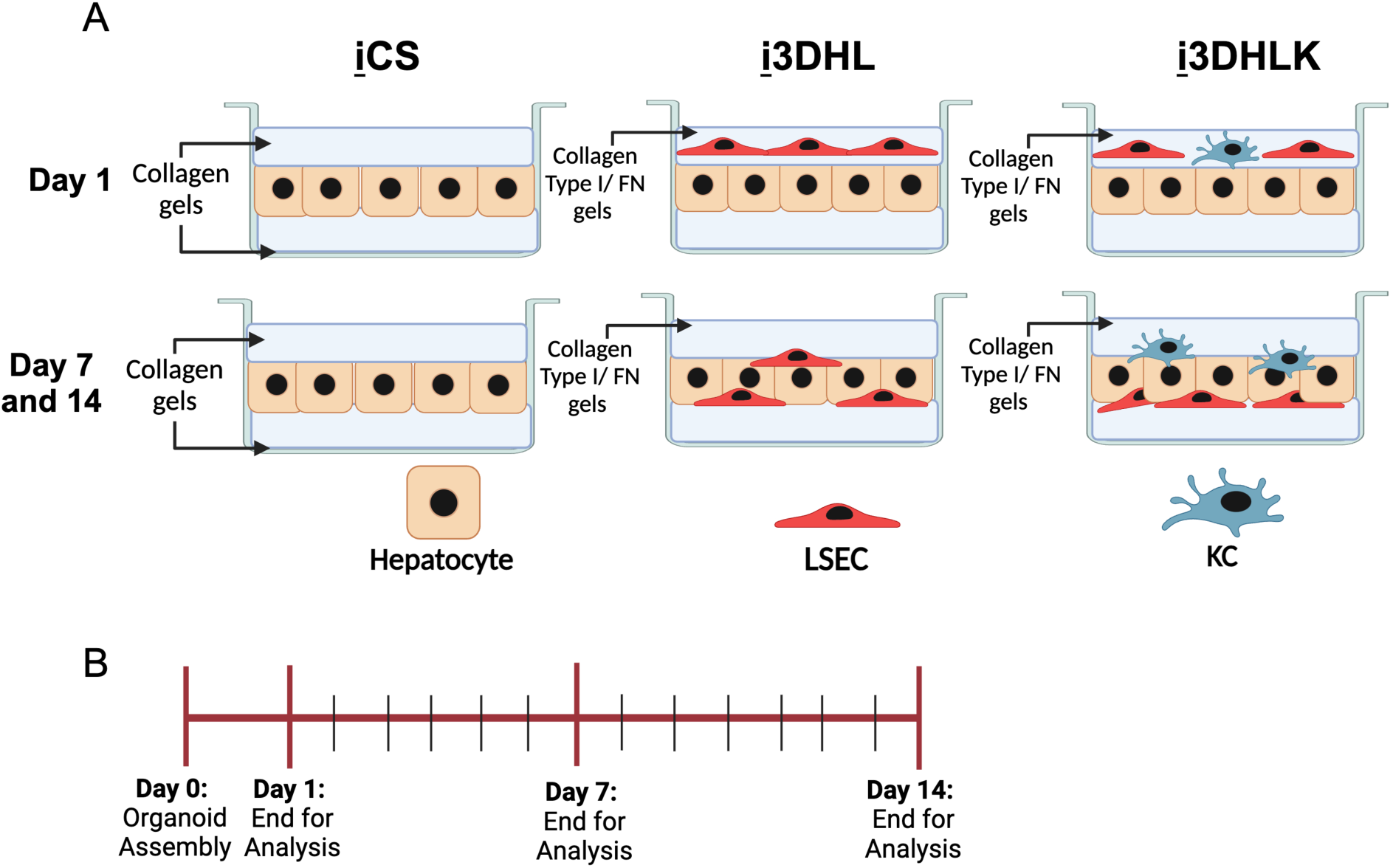
(**A**) Representative schematics of CS cultures or 3D organoids with iHLCs (**i**) (H), LSECs (L), and KCs (K) on Day 1 and Days 7 or 14. (**B**) Timeline of organoid assembly with iHLCs and PHHs.

**Figure S2.**
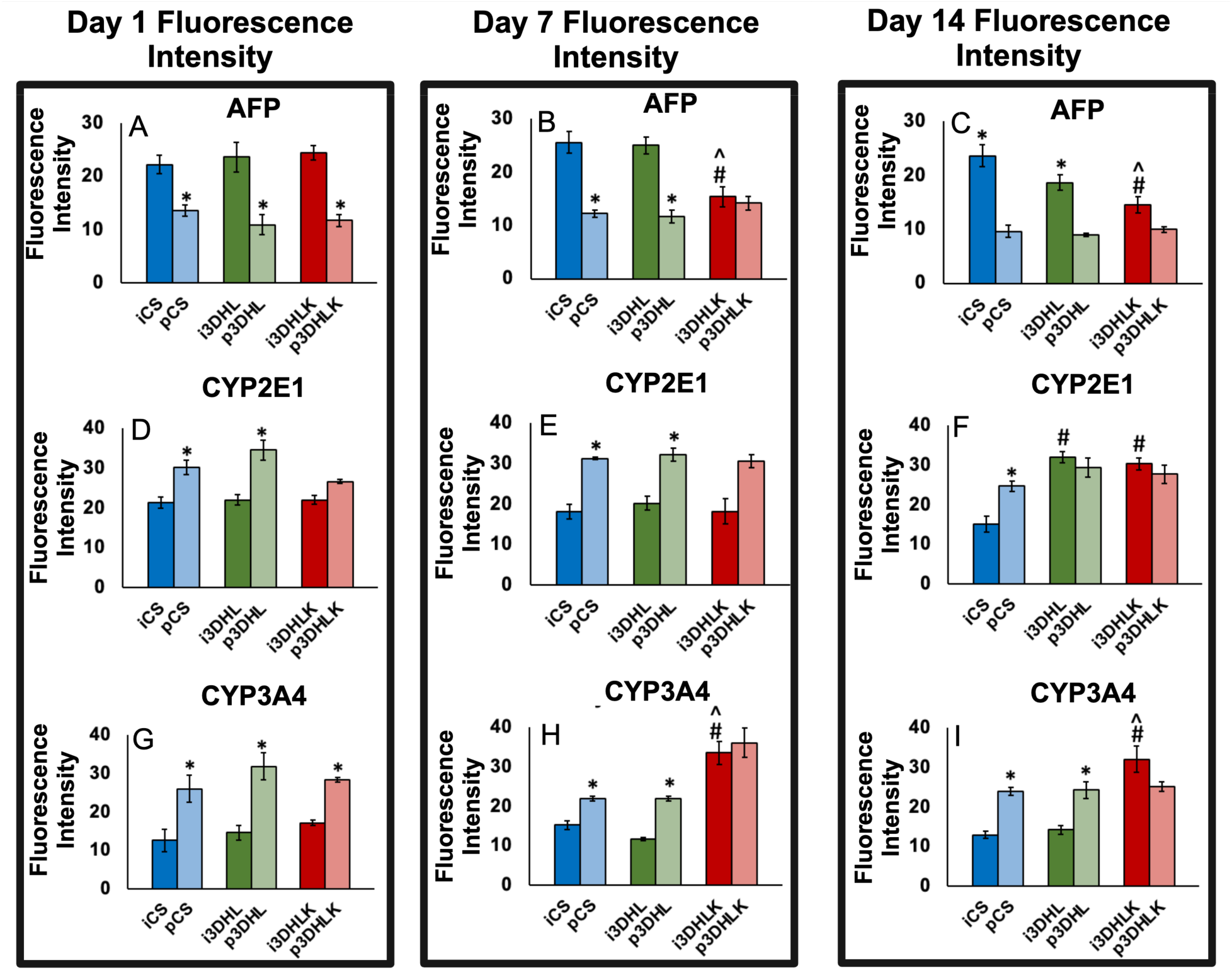
Fluorescence intensity analysis for AFP on Days 1 – 14 (**A-C**). Fluorescence intensity analysis for CYP2E1 on Days 1 – 14 (**D-F**). Fluorescence intensity analysis for CYP3A4 on Days 1 – 14 (**G-I**). \**p* < 0.05 indicates statistical significance compared to equivalent iHLC culture/ organoid. ^#^*p* < 0.05 indicates statistical significance compared to CS culture. ^*p* < 0.05 indicates statistical significance compared to 3DHL model. n = 3 biological replicates. Image analysis was conducted on ≥12 images with ≥550 cells for each organoid.

**Figure S3.**
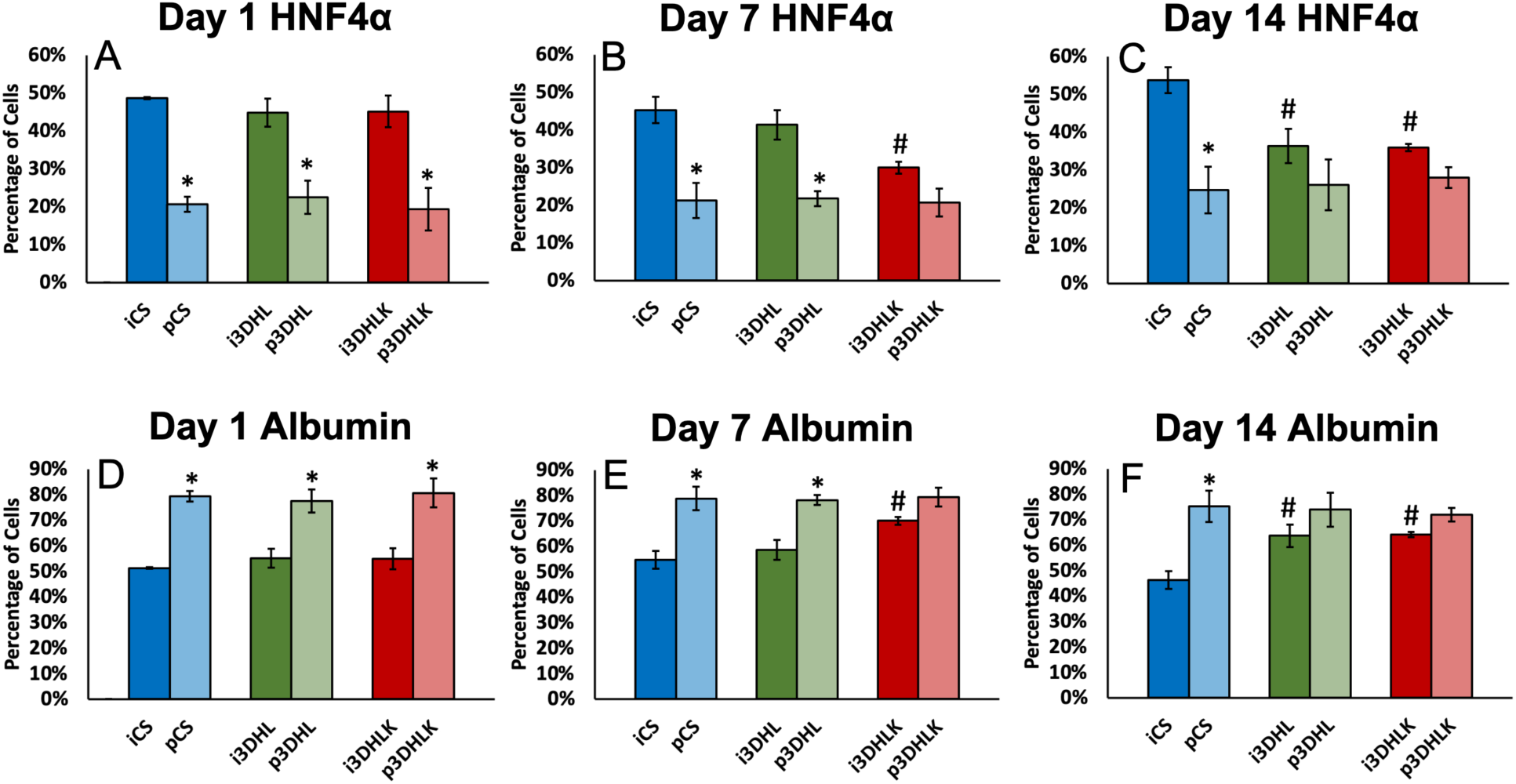
Percentage of cells expressing HNF4𝛼 on Days 1 – 14 (**A-C**). Percentage of cells expressing albumin on Days 1 – 14 (**D-F**). n = 3 biological replicates. Image analysis conducted on ≥12 images with ≥ 550 cells for each culture. \**p* < 0.05 indicates statistical significance compared to equivalent iHLC culture/ organoid. ^#^*p* < 0.05 indicates statistical significance compared to CS culture. ^*p* < 0.05 indicates statistical significance compared to 3DHL model.

**Figure S4.**
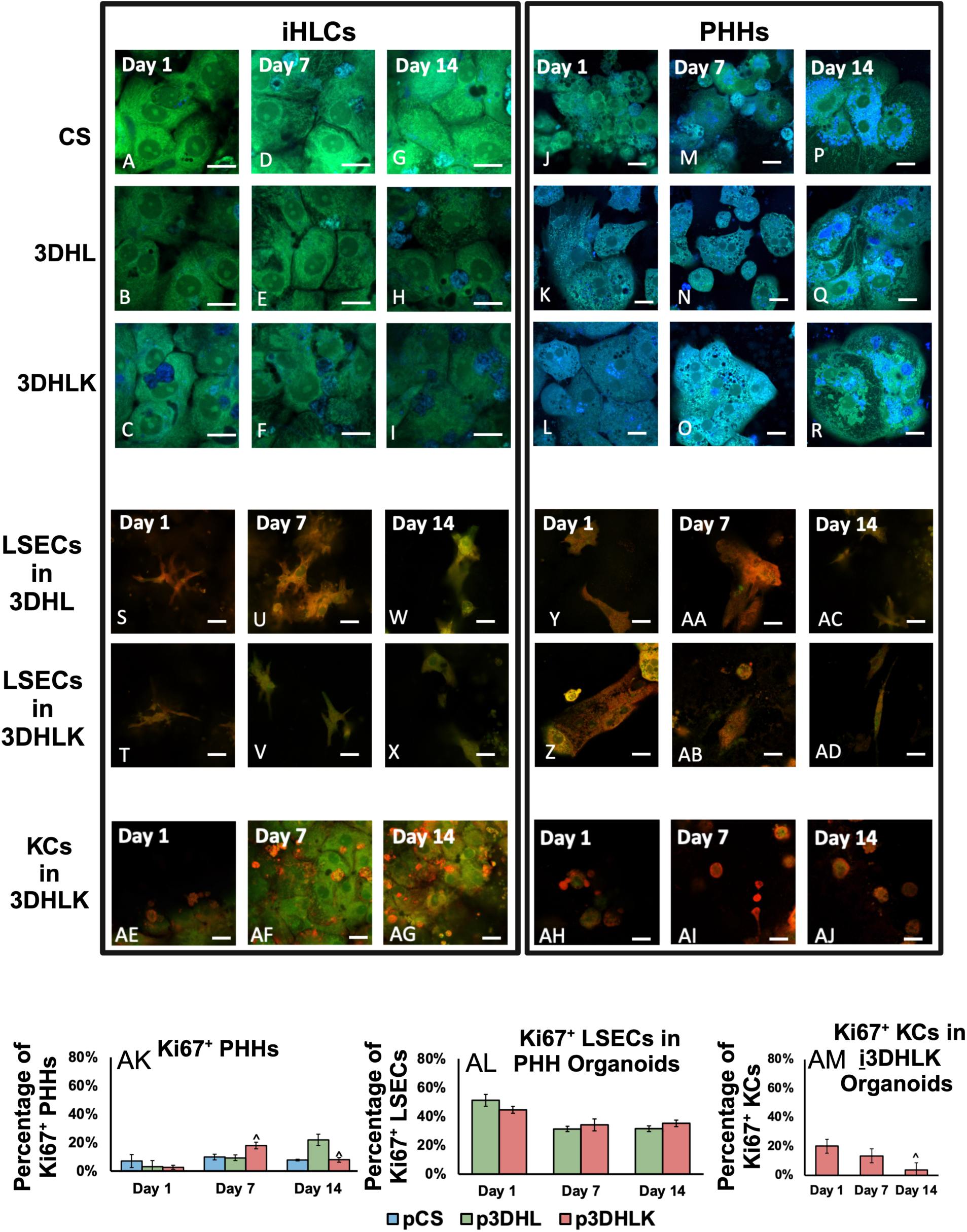
Immunostaining for Ki67 (green) and albumin (blue) for hepatocytes in iHLC (**A-I**) and PHH (**J-R**) cultures on Days 1, 7, and 14. Immunostaining for Ki67 (green) and SE-1 (red) for LSECs in iHLC (**S-X**) and PHH (**Y-AD**) organoids on Days 1, 7, and 14. Immunostaining for Ki67 (green) and CD163 (red) for KCs in **i**3DHLK (**AE-AG**) and **<u>p</u>**3DHLK (**AH-AJ**) organoids on Days 1, 7, and 14. All scale bars = 20 𝜇m. Percentages of proliferating cells in PHH organoids from Day 1 through 14. (**AK**) Percentage of Ki67^+^ PHHs on Days 1 – 14. (**AL**) Percentage of Ki67^+^ LSECs in PHH organoids on Days 1 – 14. (**AM**) Percentage of Ki67^+^ KCs in PHH organoids on Days 1 – 14. n = 3 biological replicates. Image analysis conducted on ≥12 images for each culture with ≥ 250 hepatocytes, ≥ 70 LSECs, and ≥ 100 KCs. ^*p* < 0.05 indicates statistical significance compared to Day 1.

