## Supplementary Information for "Multi-Cellular Human Liver Organoids Enable Complete Maturation of Induced Pluripotent Hepatocyte-like Cells Through Purely Endogenous Signals"

### Tables:

**Table S1.** Primary and secondary antibodies used for immunostaining phenotypic hepatic markers.

| **Primary Antibody** | **Secondary Antibody** |
|---|---|
| Rabbit anti-human HNF-4$\alpha$ (Abcam, Cambridge, UK) | Chicken anti-rabbit FITC (Novus Biologicals, Englewood, CO) |
| Sheep Serum anti-human albumin (Abcam) | Donkey anti-sheep DAPI-conjugated (Thermofisher Scientific) |
| Mouse anti-human AFP (Santa Cruz Biotechnology, Dallas, TX) | Goat anti-mouse Alexa Fluor 594 (Thermofisher Scientific) |
| Rabbit anti-human CYP2E1 (Thermofisher Scientific) | Chicken anti-rabbit Alexa Fluor 488 (Thermofisher Scientific) |
| Rabbit anti-human CYP3A4 (Thermofisher Scientific) | Chicken anti-rabbit Alexa Fluor 488 |
| Rabbit anti-human Ki67 (Abcam) | Chicken anti-rabbit Alexa Fluor 488 |
| Mouse anti-human CD163 (Abcam) | Goat anti-mouse Alexa Fluor 594 |

**Table S2**. Z-heights of each iHLC culture/ organoid on Days 1 – 14. The z-heights were measured with n≥ 15 images across n ≥ 3 biological replicates for each condition.

| | **Day 1** | **Day 7** | **Day 14** |
|---|---|---|---|
| **iCS** | 11.5 ± 1.3 | 8.4 ± 2.7 | 9.0 ± 1.7 |
| **i3DHL** | 16.0 ± 2.2 | 15.8 ± 2.0 | 14.6 ± 1.5 |
| **i3DHLK** | 15.7 ± 1.4 | 16.7 ± 2.0 | 15.8 ± 1.6 |

**Table S3**. TGF-β1 secretion in **i**3DHLK and **p**3DHLK organoids on Days 1 and 14. Culture medium values were subtracted from the raw data and then the concentration was normalized to the protein content in the organoid at each time point (pg/ mg protein). n = 3 biological replicates. (-) indicates that the concentration was undetectable.

| **Media/Culture** | **pg/ mL** | **Day 1 (pg/ mg protein)** | **Day 14 (pg/ mg protein)** |
|---|---|---|---|
| **iHLC Media** | – | | |
| **PHH Media** | 159.0 ± 89.0 | | |
| **i3DHLK** | | 117.4 ± 179.9 | 35.8 ± 59.0 |
| **p3DHLK** | | 55.7 ± 58.9 | 55.5 ± 51.8 |

***APAP ADMINISTRATION:***

**Table S4**. Percentage of apoptotic, necrotic, and total cell death measured 24 h after 2.5 mM APAP administration on Day 1. n ≥ 3 biological replicates for each condition.

| | Untreated on Day 1 | | | Administration on Day 1 | | |
|---|---|---|---|---|---|---|
| **Model** | **Apoptosis (%)** | **Necrosis (%)** | **Cell Death (%)** | **Apoptosis (%)** | **Necrosis (%)** | **Cell Death (%)** |
| **i**CS | 5.9 ± 2.2 | 0.9 ± 0.5 | 6.7 ± 2.4 | 5.3 ± 2.8 | 4.7 ± 1.9 | 10.0 ± 1.7 |
| **i**3DHL | 2.7 ± 1.2 | 5.1 ± 1.7 | 7.8 ± 2.7 | 2.2 ± 0.6 | 4.5 ± 1.1 | 6.7 ± 0.6 |
| **i**3DHLK | 3.4 ± 1.9 | 2.1 ± 0.9 | 5.5 ± 2.9 | 3.6 ± 2.7 | 2.3 ± 1.0 | 5.8 ± 3.7 |
| **p**CS | 6.8 ± 3.5 | 3.4 ± 2.4 | 10.2 ± 4.5 | 14.3 ± 1.6 | 3.6 ± 0.6 | 17.9 ± 2.2 |
| **p**3DHL | 8.4 ± 2.9 | 4.0 ± 0.6 | 12.3 ± 3.2 | 12.1 ± 3.5 | 14.2 ± 0.7 | 26.3 ± 9.4 |
| **p**3DHLK | 3.6 ± 1.4 | 2.8 ± 0.6 | 6.8 ± 2.0 | 16.2 ± 1.4 | 25.9 ± 3.3 | 42.1 ± 3.0 |

**Table S5**. Percentage of apoptotic, necrotic, and total cell death measured in untreated samples on Days 7 and 14. n ≥ 3 biological replicates for each condition.

| | Untreated on Day 7 | | | Untreated on Day 14 | | |
|---|---|---|---|---|---|---|
| **Model** | **Apoptosis (%)** | **Necrosis (%)** | **Cell Death (%)** | **Apoptosis (%)** | **Necrosis (%)** | **Cell Death (%)** |
| **i**CS | 0.7 ± 0.2 | 6.2 ± 1.2 | 6.9 ± 1.1 | 5.4 ± 2.5 | 2.9 ± 1.2 | 8.4 ± 3.0 |
| **i**3DHL | 1.7 ± 0.8 | 4.1 ± 1.9 | 5.8 ± 2.6 | 5.0 ± 1.4 | 1.8 ± 0.5 | 6.7 ± 1.8 |
| **i**3DHLK | 2.5 ± 1.1 | 1.3 ± 1.4 | 3.9 ± 1.3 | 4.0 ± 2.03 | 0.7 ± 0.7 | 4.7 ± 2.5 |
| **p**CS | 6.4 ± 3.5 | 5.0 ± 2.2 | 11.4 ± 5.5 | 9.5 ± 4.7 | 14.6 ± 3.5 | 24.1 ± 7.4 |
| **p**3DHL | 7.8 ± 2.9 | 5.8 ± 1.6 | 13.6 ± 4.1 | 7.8 ± 2.1 | 19.1 ± 2.6 | 26.9 ± 4.5 |
| **p**3DHLK | 3.7 ± 1.5 | 4.0 ± 1.2 | 7.7 ± 0.5 | 8.5 ± 2.3 | 17.7 ± 3.2 | 26.1 ± 3.6 |

***EtOH ADMINISTRATION:***

**Table S6**. Percentage of apoptotic, necrotic, and total cell death measured 24 h after 160 mM EtOH administration on Day 1. n ≥ 3 biological replicates for each condition.

| | Untreated on Day 1 | | | Administration on Day 1 | | |
|---|---|---|---|---|---|---|
| **Model** | **Apoptosis (%)** | **Necrosis (%)** | **Cell Death (%)** | **Apoptosis (%)** | **Necrosis (%)** | **Cell Death (%)** |
| **i**CS | 5.9 ± 2.2 | 0.9 ± 0.5 | 6.7 ± 2.4 | 3.8 ± 1.4 | 1.4 ± 0.6 | 5.2 ± 1.9 |
| **i**3DHL | 2.7 ± 1.2 | 5.1 ± 1.7 | 7.8 ± 2.7 | 3.9 ± 3.2 | 5.2 ± 1.7 | 9.1 ± 1.6 |
| **i**3DHLK | 3.4 ± 1.9 | 2.1 ± 0.9 | 5.5 ± 2.9 | 3.5 ± 1.2 | 2.3 ± 0.9 | 5.8 ± 1.6 |
| **p**CS | 6.8 ± 3.5 | 3.4 ± 2.4 | 10.2 ± 4.5 | 13.3 ± 3.0 | 14.0 ± 1.0 | 27.3 ± 2.9 |
| **p**3DHL | 8.4 ± 2.9 | 4.0 ± 0.6 | 12.3 ± 3.2 | 17.0 ± 5.5 | 15.9 ± 4.2 | 32.9 ± 7.6 |
| **p**3DHLK | 3.6 ± 1.4 | 2.8 ± 0.6 | 6.8 ± 2.0 | 16.5 ± 5.0 | 24.8 ± 1.4 | 41.3 ± 4.3 |

**Table S7**. Percentage of apoptotic, necrotic, and total cell death measured in untreated samples on Days 7 and 14. n ≥ 3 biological replicates for each condition.

| | Untreated on Day 7 | | | Untreated on Day 14 | | |
|---|---|---|---|---|---|---|
| **Model** | **Apoptosis (%)** | **Necrosis (%)** | **Cell Death (%)** | **Apoptosis (%)** | **Necrosis (%)** | **Cell Death (%)** |
| **i**CS | 2.6 ± 1.3 | 2.8 ± 1.4 | 5.4 ± 2.4 | 6.8 ± 2.5 | 5.6 ± 3.1 | 12.4 ± 5.6 |
| **i**3DHL | 4.4 ± 2.5 | 0.8 ± 0.8 | 5.2 ± 3.3 | 2.2 ± 1.7 | 1.7 ± 0.6 | 3.9 ± 1.7 |
| **i**3DHLK | 2.0 ± 0.1 | 0.5 ± 0.3 | 2.4 ± 0.4 | 5.0 ± 0.9 | 2.5 ± 0.5 | 7.5 ± 1.4 |
| **p**CS | 6.4 ± 3.5 | 5.0 ± 2.2 | 11.4 ± 5.5 | 9.5 ± 4.7 | 14.6 ± 3.5 | 24.1 ± 7.4 |
| **p**3DHL | 7.8 ± 2.9 | 5.8 ± 1.6 | 13.6 ± 4.1 | 7.8 ± 2.1 | 19.1 ± 2.6 | 26.9 ± 4.5 |
| **p**3DHLK | 3.7 ± 1.5 | 4.0 ± 1.2 | 7.7 ± 0.5 | 8.5 ± 2.3 | 17.7 ± 3.2 | 26.1 ± 3.6 |

***RIF ADMINISTRATION:***

**Table S8**. Percentage of apoptotic, necrotic, and total cell death measured 24 h after 0.36 $\mu$M RIF administration on Day 1. n ≥ 3 biological replicates for each condition.

| | **Untreated on Day 1** | | | **Administration on Day 1** | | |
|---|---|---|---|---|---|---|
| **Model** | **Apoptosis (%)** | **Necrosis (%)** | **Cell Death (%)** | **Apoptosis (%)** | **Necrosis (%)** | **Cell Death (%)** |
| iCS | 6.6 ± 1.4 | 1.4 ± 0.6 | 8.0 ± 1.5 | 7.7 ± 4.0 | 2.7 ± 1.6 | 10.4 ± 5.6 |
| i3DHL | 7.0 ± 1.7 | 1.5 ± 0.2 | 8.5 ± 1.7 | 3.6 ± 1.8 | 2.5 ± 1.2 | 6.1 ± 3.0 |
| i3DHLK | 3.4 ± 1.2 | 1.8 ± 1.3 | 5.1 ± 2.4 | 5.2 ± 3.0 | 2.7 ± 1.4 | 7.9 ± 1.9 |
| pCS | 3.4 ± 2.3 | 0.7 ± 0.6 | 4.1 ± 2.0 | 5.7 ± 1.9 | 3.9 ± 2.8 | 9.6 ± 4.7 |
| p3DHL | 3.0 ± 2.0 | 2.8 ± 1.1 | 5.9 ± 2.9 | 9.2 ± 0.8 | 7.4 ± 0.9 | 16.5 ± 0.4 |
| p3DHLK | 4.1 ± 1.0 | 1.5 ± 1.3 | 5.6 ± 0.3 | 21.8 ± 2.6 | 19.3 ± 1.8 | 41.1 ± 1.2 |

**Table S9**. Percentage of apoptotic, necrotic, and total cell death measured in untreated samples on Days 7 and 14. n ≥ 3 biological replicates for each condition.

| | **Untreated on Day 7** | | | **Untreated on Day 14** | | |
|---|---|---|---|---|---|---|
| **Model** | **Apoptosis (%)** | **Necrosis (%)** | **Cell Death (%)** | **Apoptosis (%)** | **Necrosis (%)** | **Cell Death (%)** |
| iCS | 5.1 ± 1.0 | 2.4 ± 0.6 | 7.5 ± 1.5 | 2.9 ± 1.1 | 4.0 ± 1.7 | 6.8 ± 2.7 |
| i3DHL | 5.1 ± 0.7 | 1.6 ± 0.5 | 6.7 ± 0.3 | 4.3 ± 1.0 | 3.2 ± 2.1 | 7.5 ± 2.2 |
| i3DHLK | 4.5 ± 0.8 | 2.2 ± 2.5 | 6.7 ± 2.7 | 4.0 ± 1.2 | 2.6 ± 1.0 | 6.6 ± 0.8 |
| pCS | 5.8 ± 2.7 | 0.8 ± 0.8 | 6.6 ± 2.7 | 4.3 ± 0.5 | 3.4 ± 1.8 | 7.7 ± 2.3 |
| p3DHL | 4.0 ± 1.1 | 4.4 ± 3.1 | 8.3 ± 2.8 | 6.7 ± 0.1 | 3.4 ± 1.10 | 10.1 ± 1.2 |
| p3DHLK | 6.5 ± 1.0 | 3.1 ± 1.7 | 9.6 ± 2.1 | 5.8 ± 0.8 | 3.5 ± 0.9 | 9.3 ± 0.2 |

**Table S10**. HGF secretion in iHLC and PHH organoids/cultures and LSEC and KC monocultures on Days 1, 7, and 14. Culture medium values were subtracted from the raw data and then the concentration was normalized to the protein content in the organoid/culture at each time point (pg/ mg protein). n = 3 biological replicates.

| **Media/Culture** | **pg/ mL** | **Day 1 (pg/ mg protein)** | **Day 7 (pg/ mg protein)** | **Day 14 (pg/ mg protein)** |
|---|---|---|---|---|
| **iHLC Media** | 46.0 ± 14.0 | | | |
| **PHH Media** | 353.0 ± 42.0 | | | |
| **LSEC Monoculture** | | 302.3 ± 40.3 | | |
| **KC Monoculture** | | 2.2 ± 1.9 | | |
| **iCS** | | 35.0 ± 8.5 | 39.7 ± 5.8 | 33.0 ± 5.3 |
| **i3DHL** | | 39.3 ± 15.0 | 115.6 ± 33.7 | 123.3 ± 43.9 |
| **i3DHLK** | | 44.5 ± 8.2 | 76.4 ± 2.9 | 76.0 ± 14.1 |
| **pCS** | | – | – | – |
| **p3DHL** | | – | – | – |
| **p3DHLK** | | – | – | – |

**Table S11**. OSM secretion in iHLC and PHH organoids/cultures and LSEC and KC monocultures on Days 1, 7, and 14. Culture medium values were subtracted from the raw data and then the concentration was normalized to the protein content in the organoid/culture at each time point (pg/ mg protein). n = 3 biological replicates.

| Media/Culture | pg/ mL | Day 1 (pg/ mg protein) | Day 7 (pg/ mg protein) | Day 14 (pg/ mg protein) |
|---|---|---|---|---|
| **iHLC Media** | 11.3 ± 0.6 | | | |
| **PHH Media** | 42.7 ± 7.6 | | | |
| **LSEC Monoculture** | | 0.4 ± 0.7 | | |
| **KC Monoculture** | | 12.9 ± 4.6 | | |
| **iCS** | | 2.8 ± 0.8 | 4.1 ± 1.1 | 3.0 ± 0.3 |
| **i3DHL** | | 2.2 ± 0.5 | 2.7 ± 0.8 | 4.1 ± 1.2 |
| **i3DHLK** | | 4.9 ± 2.2 | 4.1 ± 1.2 | 12.4 ± 1.0 |
| **pCS** | | – | – | – |
| **p3DHL** | | – | – | – |
| **p3DHLK** | | – | – | – |

**Table S12**. PGE2 secretion in iHLC and PHH organoids/cultures and LSEC and KC monocultures on Days 1, 7, and 14. Culture medium values were subtracted from the raw data and then the concentration was normalized to the protein content in the organoid/culture at each time point (pg/ mg protein). n = 3 biological replicates.

| Media/Culture | pg/ mL | Day 1 (pg/ mg protein) | Day 7 (pg/ mg protein) | Day 14 (pg/ mg protein) |
|---|---|---|---|---|
| **iHLC Media** | 46.3 ± 6.0 | | | |
| **PHH Media** | 34.0 ± 2.6 | | | |
| **LSEC Monoculture** | | 15.4 ± 0.8 | | |
| **KC Monoculture** | | 7.0 ± 6.1 | | |
| **iCS** | | 6.0 ± 7.3 | 7.8 ± 11.7 | 10.8 ± 2.3 |
| **i3DHL** | | 10.8 ± 10.2 | 5.3 ± 8.0 | 17.0 ± 6.0 |
| **i3DHLK** | | 39.4 ± 6.7 | 8.1 ± 4.5 | 6.1 ± 4.6 |
| **pCS** | | 14.8 ± 3.8 | 30.4 ± 12.3 | 43.1 ± 5.0 |
| **p3DHL** | | 17.6 ± 9.4 | 32.3 ± 9.1 | 96.4 ± 11.8 |
| **p3DHLK** | | 38.4 ± 18.3 | 73.6 ± 2.3 | 13.0 ± 9.8 |

**Table S13**. Albumin secretion in iHLC and PHH organoids/cultures and LSEC and KC monocultures on Days 1, 7, and 14. Culture medium values were subtracted from the raw data and then the concentration was normalized to the protein content in the organoid/culture at each time point (pg/ mg protein). n = 3 biological replicates.

| Media/Culture | pg/ mL | Day 1 | Day 7 | Day 14 |
|---|---|---|---|---|

| | | (pg/ mg protein) | (pg/ mg protein) | (pg/ mg protein) |
|---|---|---|---|---|
| **iHLC Media** | 9169 ± 693 | | | |
| **PHH Media** | 7448 ± 117 | | | |
| **LSEC Monoculture** | | 52.8 ± 91.41 | | |
| **KC Monoculture** | | 376.4 ± 651.9 | | |
| **iCS** | | $17.4 \times 10^5 \pm 0.6 \times 10^5$ | $19.9 \times 10^5 \pm 1.3 \times 10^5$ | $13.1 \times 10^5 \pm 1.3 \times 10^5$ |
| **i3DHL** | | $31.7 \times 10^5 \pm 1.6 \times 10^5$ | $35.2 \times 10^5 \pm 1.8 \times 10^5$ | $30.9 \times 10^5 \pm 4.0 \times 10^5$ |
| **i3DHLK** | | $34.0 \times 10^5 \pm 0.1 \times 10^5$ | $33.5 \times 10^5 \pm 1.2 \times 10^5$ | $24.2 \times 10^5 \pm 9.1 \times 10^5$ |
| **pCS** | | $4.6 \times 10^5 \pm 0.7 \times 10^5$ | $5.3 \times 10^5 \pm 3.0 \times 10^5$ | $7.3 \times 10^5 \pm 2.5 \times 10^5$ |
| **p3DHL** | | $4.2 \times 10^5 \pm 0.6 \times 10^5$ | $15.31 \times 10^5 \pm 1.57 \times 10^5$ | $25.5 \times 10^5 \pm 1.6 \times 10^5$ |
| **p3DHLK** | | $4.6 \times 10^5 \pm 0.4 \times 10^5$ | $16.7 \times 10^5 \pm 3.6 \times 10^5$ | $23.9 \times 10^5 \pm 0.6 \times 10^5$ |

Movies:

S1. 3D rendering of iCS cultures on Day 1.

S2. 3D rendering of i3DHL organoids on Day 1.

S3. 3D rendering of i3DHLK organoids on Day 1.

S4. 3D rendering of iCS cultures on Day 14.

S5. 3D rendering of i3DHL organoids on Day 14.

S6. 3D rendering of i3DHLK organoids on Day 14.
